# Warming up to a new coat: moulting king penguins exhibit hyperthermia and increased peripheral heat loss

**DOI:** 10.1101/2025.03.25.645083

**Authors:** Juan D Zuluaga, Emmanuel Pretti, Aude Leynaert, Elsa Marçon, Antoine Stier, Agnès Lewden

## Abstract

Penguins are among the most specialized thermoregulators on the planet, however, the same adaptations that maximize heat retention underwater likely hinder heat dissipation on land, possibly creating dangerous thermoregulatory challenges when encountering warming terrestrial habitats. Penguins are subject to strictly terrestrial phases, such as moulting, when metabolic heat production, insulation, and energetic constraints are heightened. We assessed thermoregulation in moulting captive king penguins (*Aptenodytes patagonicus*) using simultaneous measurements of core and surface temperatures to test two hypotheses. Under the *thermal challenge* hypothesis, an initial rise in heat dissipation effort (i.e., increased peripheral vasomotion) followed by a rise in core temperature would indicate failure to prevent hyperthermia. Under the *warm-up* hypothesis, an initial rise of core temperature concomitant or followed by an increase of peripheral vasomotion would indicate regulated hyperthermia, possibly to accelerate feather development. Core and surface temperatures increased drastically but concomitantly during moult, providing tentative support for the *warm-up* hypothesis. Moulting penguins did not pant, suggesting that peripheral heat dissipation was sufficient to regulate moulting-induced hyperthermia. Core and subcutaneous temperatures in wild individuals resembled patterns measured in captivity, despite lower heat load and additional options for behavioural thermoregulation. These results indicate that hyperthermia is prevalent in moulting king penguins, and documenting the timing of temperature changes provides novel insights for the moulting physiology of penguins. Because moulting-induced hyperthermia may contribute to heat load, we caution that moulting may increase the susceptibility of wild penguins to heat stress, especially as regions near the poles warm at a disproportionately rapid rate.

## Introduction

Maintaining thermal insulation is essential for endotherms to thrive in their natural environments (1). Fur and feathers contribute to thermal insulation in most mammals and birds, but these keratinaceous structures degrade through wear and breakage as they age. Thus, the maintenance of proper insulation from fur or feathers through time requires their replacement through shedding: a protracted, year-round replacement, or moulting: a contracted, punctuated replacement (1). Feathers are unique among morphological traits due to the variety of functions they serve, with major roles in crypsis, mate choice, species signalling, thermoregulation, waterproofing, flight, and protection (2). Moulting is thus considered a key life-history stage because it conditions the ability of individuals to survive and reproduce in their environment (3).

Moult has been well characterized in various avian models from a metabolic perspective. Moult requires a massive amount of protein because feathers are composed of approximately 90% protein and can represent or even exceed 10% of body mass and 25% of total body protein content (4). Synthesizing new feathers also requires an increase in energy expenditure, with moulting birds generally exhibiting a metabolic rate *ca.* 9-111% higher than non-moulting individuals (4). From a thermoregulatory perspective, moult has been less well characterized to the best of our knowledge. The few studies available in birds suggest that an increase of *ca.* 20-60% of thermal conductance occurs during moult (5, 6), which may be explained by a decrease in insulation and/or an increase in evaporative heat transfer due to increased body-water turnover (4). Increases in thermal conductance have been shown to raise the lower critical temperature in long-eared owls (*Asio otus*) from approximately 18°C to 24°C (7) and in emperor penguins (*Aptenodytes forsteri*) from approximately −10°C to 0°C (8). Thus, moult has mostly been considered thermally challenging under cold environmental conditions. Alternatively, it has been hypothesized that the increased metabolism of moulting birds could more than compensate for the decrease in insulation, and thus that the increased heat generation may pose a challenge for moulting birds (4, 9, 10).

Penguins have specialized vascular arrangements that provide countercurrent heat exchange in the legs (11), head (12), and wings (13). Penguins also boast efficient insulation through a layer of subcutaneous fat (14), and specialized plumage comprised of an insulating underlayer and a waterproof overlayer (8, 15). Despite penguins’ adaptations for heat retention underwater, they must breed and moult on land, where dissipating heat is more difficult than in water (thermal conductance is 2.2-4.8 times higher in water than in air; (16)). Penguins undergo a catastrophic moult once a year, replacing their entire plumage extremely quickly (between 13 and 34 days; (17)), which involves an increase of metabolic rate (and thus of heat production) of *ca.* 30-50% (18, 19). Catastrophic moult involves the temporary overlap of new and old feather layers (20), which increases thermal insulation (10). Penguins seem relatively sensitive to heat stress while on land (21–23), and climate change is pronounced in polar and sub-polar regions (24, 25), where many penguin species moult on land during the warmest season (26). Thus, if heat dissipation is indeed a challenge during moult (4, 9, 10), the increasing temperatures penguins are facing while moulting on land may challenge these iconic species.

Captive Gentoo penguins held within their thermoneutral zone had increased thermal insulation and heat dissipation effort during the overlap of old and new feather layers (10). Therefore, moulting seems to represent a challenge for heat dissipation in Gentoo penguins, but the efficiency or inefficiency of their thermoregulatory response remains unknown because core body temperature was not measured. We define the *thermal challenge* hypothesis as the scenario where core body temperature rises because of failure to dissipate heat. Alternatively, it has been suggested that birds may elevate core body temperature while moulting, hypothetically under physiological control, to favour feather growth (3), an idea we are referring to as the *warm-up* hypothesis. If heat dissipation effort increases but core temperature does not, this would indicate that heat dissipation capacity is sufficient to offset the increased metabolism (4) and insulation (10) that are known to occur during moult. If heat dissipation effort increases and body temperature increases afterwards, this would indicate a failure to regulate body temperature, thus supporting the *thermal challenge* hypothesis.

To test these hypotheses, we simultaneously recorded panting behaviour, heat dissipation effort (*i.e.* surface temperature of thermal windows such as flippers, bill and feet), insulation (*i.e.* surface temperature of the trunk) and core body temperature in captive king penguins before, during and after the moult. To address ecological relevance of the results obtained in captivity, we also include behavioural and physiological data of wild king penguins, although limited in sample size. Under the thermal challenge hypothesis, we predict that wild king penguins will be less likely to exhibit hyperthermia because environmental conditions are colder in the field and because wild king penguins have additional methods to cool themselves (*e.g.,* immersing feet in cold water). If core body temperature increases before heat dissipation effort, or if both increase simultaneously, this would indicate a regulated increase in core body temperature, thus supporting the *warm-up* hypothesis. Under the *warm-up* hypothesis, we predict that both captive and wild king penguins will exhibit hyperthermia despite environmental and behavioural variation.

## Materials and Methods

### Captive king penguins

We studied 8 captive adult king penguins (*Aptenodytes patagonicus*) in the spring of 2024 at Océanopolis aquarium, Brest, France. The birds were individually identified by their coloured plastic band, and six individuals moulted during the study period. Individuals were maintained indoors at 8.12 ± 0.07°C (mean ± SE), well within their estimated thermoneutral zone (*i.e.* L-5 toL+22 °C; (27)). They had permanent access to free water to swim/dive, but no possibility to only immerse their feet. Researchers were the first to enter the enclosure after a full night with no disturbance to collect thermal images shortly before 8:00 AM, and keepers entered afterwards throughout the day for regular enclosure cleanings, feeding non-moulting birds (*i.e.* moulting birds are naturally fasting and stay on land even in captivity). The lighting program used in the enclosure includes a monthly variation in artificial light, with exposure varying between 11 and 13 h of light per day during the study. Following the 7-stage moult categorization (10), we assigned moult stage each day for each individual from uniform old plumage (M1) to uniform new plumage (M7) before surface temperature data extraction. To investigate the continuous changes that occur within the visibly distinct moult stages, we tracked the moulting process numerically by assigning a moult day for each bird, which began at day 0 when we visually detected changes in the plumage structure (*i.e.* the start of M2, (10)). Moult actually starts before any visual changes occur (*i.e.* feather growth below the skin), which occurs at sea in wild penguins and has been estimated to last *ca.* 10 days (20).

Individuals were fed with an ingestible core temperature (T_core_) data logger (Bodycap, Anipill, France, accuracy ± 0.2°C between 25 and 45°C, resolution ± 0.01°C, sampling rate = 5min) hidden in their normal diet (*i.e.* without handling) when they began exhibiting hyperphagia, which precedes the fasting period during catastrophic moult (28). Individuals were equipped at this stage because retention time in the stomach is quite variable (*ca.* 7 to 80 days in wild king penguins, *pers. obs.*). We expect that stomach temperature provides a close approximation of core body temperature during the moult because temperature measurements in fasting penguins are not influenced by food intake. To avoid collecting measurements of core temperature that may be influenced by food intake in non-moulting individuals, we collected temperature data of all birds in the mornings shortly before 8:00 AM after a full fasting night (*i.e.* 30 minutes after lights turned on, and before any keepers had entered the enclosure to feed the non-moulting birds). Five moulting individuals were successfully equipped before the onset of visible plumage change, and one at the time of early plumage change (*i.e.* early M2 phase, (10)). Only one (non-moulting) individual lost the logger before the completion of the study, 32 days after ingestion. Moulting individuals were equipped on average 16.5 ± 4.8 days before visible changes in plumage occurred. Data were downloaded remotely with an Anilogger® monitor placed inside the enclosure throughout the study.

Upon entering the enclosure to collect data, the presence/absence of panting behaviour was recorded for each individual (22), and we used a thermal imaging camera (three different cameras were used due to technical issues: E96, E8, or T640; resolutions provided in Electronic Supplementary Materials, *ESM*) to collect images of each individual from a distance of 1 m (10). We collected images of birds in the profile orientation (29). Air temperature (T_a_) and relative humidity (RH) inside the enclosure were measured using a weather station (Kestrel 5400; Kestrel® instrument, United Kingdom). Thermal imaging collection started on average 23.3 ± 5.4 days before visible changes in plumage occurred. Pictures of individuals being wet at the time of thermal imaging were excluded from the analysis (*i.e.* n = 36 images from non-moulting individuals, or moulting ones with moult day > 20). We report the average core body temperature from 7:30 to 8:00 AM. This approach resulted in a sample size of 291 datapoints with body surface and core temperatures measured simultaneously.

After collecting thermal images, we extracted surface temperatures using FLIR Thermal Studio Pro. We set emissivity to 0.98 (10) and we specified T_a_ and RH for each image using measurements collected during each thermal imaging session. We then measured the average surface temperatures of the bill, flipper, and foot (T_bill_, T_flipper_, and T_foot_, respectively) by drawing separate region of interest (ROI) polygons for each area. We measured the maximum surface temperature of the periorbital region (henceforth T_eye_; (30)) using an ellipse. We measured ground temperature (T_ground_) below the bird using a line segment the length of the foot. To determine a representative trunk surface temperature (T_trunk_) despite irregular feather loss, we measured the temperature of both the old and newly grown plumage (T_old_ and T_new_) using a square with side length equal to ∼one third the length of the flipper. We used visual landmarks to measure T_ground_, T_old_, and T_new_ because image resolution varied between the three cameras. After standardizing between the cameras (see *ESM*), we used values of T_old_ and T_new_ to calculate T_trunk_ according to (10). The moulting pattern of king penguins is sufficiently patchy for squares to be consistently drawn in a section of completely old or new plumage with no overlap.

Data analysis was conducted in R Version 4.3.3 (31). We first calculated wet-bulb temperature (T_w_) in R package HeatStress (32) using the measurements of T_a_ and RH collected while sampling. Following guidance from two external (*i.e.,* non-author) experts in the field of thermography, we standardized measurements made with the three cameras using two approaches (*i.e.* see ESM S0 for details). We then modelled T_core_, T_eye_, T_bill_, T_flipper_, T_foot_, and T_trunk_ as a function of moult day. To capture the non-linear changes of body temperatures over the course of the moult (in moulting birds only), we created segmented models based on calculated changes in slope (*i.e.,* breakpoints; (33)). We determined the presence/timing of breakpoints and the associated 95% intervals using the R package segmented (34). After identifying breakpoints, we fitted separate linear mixed models ((lmer()) in R package lme4 (35)) for each time period as determined by the breakpoints for moulting birds. Although thermal conditions were held relatively constant (Fig. S1), we included T_w_ as a fixed effect in all global models to account for any subtle variation in thermal conditions; we used T_ground_ instead of T_w_ for the models of foot temperature according to (10). Non-moulting birds were analysed in separate mixed models and were assigned an arbitrary moulting day for graphical purpose only (Fig. 1). We included the bird’s assigned identification number (bird ID) as a random intercept in all models to account for repeated measures. Model support was determined using AICc (36) using the function dredge() in R package MuMIn (37). We report the top-ranked model with parameter estimates for moult day and T_w_ or T_ground_ (Table S1). We assessed the fit of the top models by calculating the conditional Nakagawa’s R^2^ using function r.squaredGLMM() in R package MuMIn. We visually represented findings from our models by plotting model predictions obtained from package ggeffects (38) along with raw data for each response variable using package ggplot2 (39). Finally, predicted values and the associated 95% confidence intervals were extracted from key stages based on breakpoint (*i.e.* pre-moult, peak-moult, post-moult) as well as non-moulting individuals using semi-parametric bootstrapping (n = 1000 simulations) with confint() and bootMer() functions of R package lme4 (35)).

**Figure 1.**
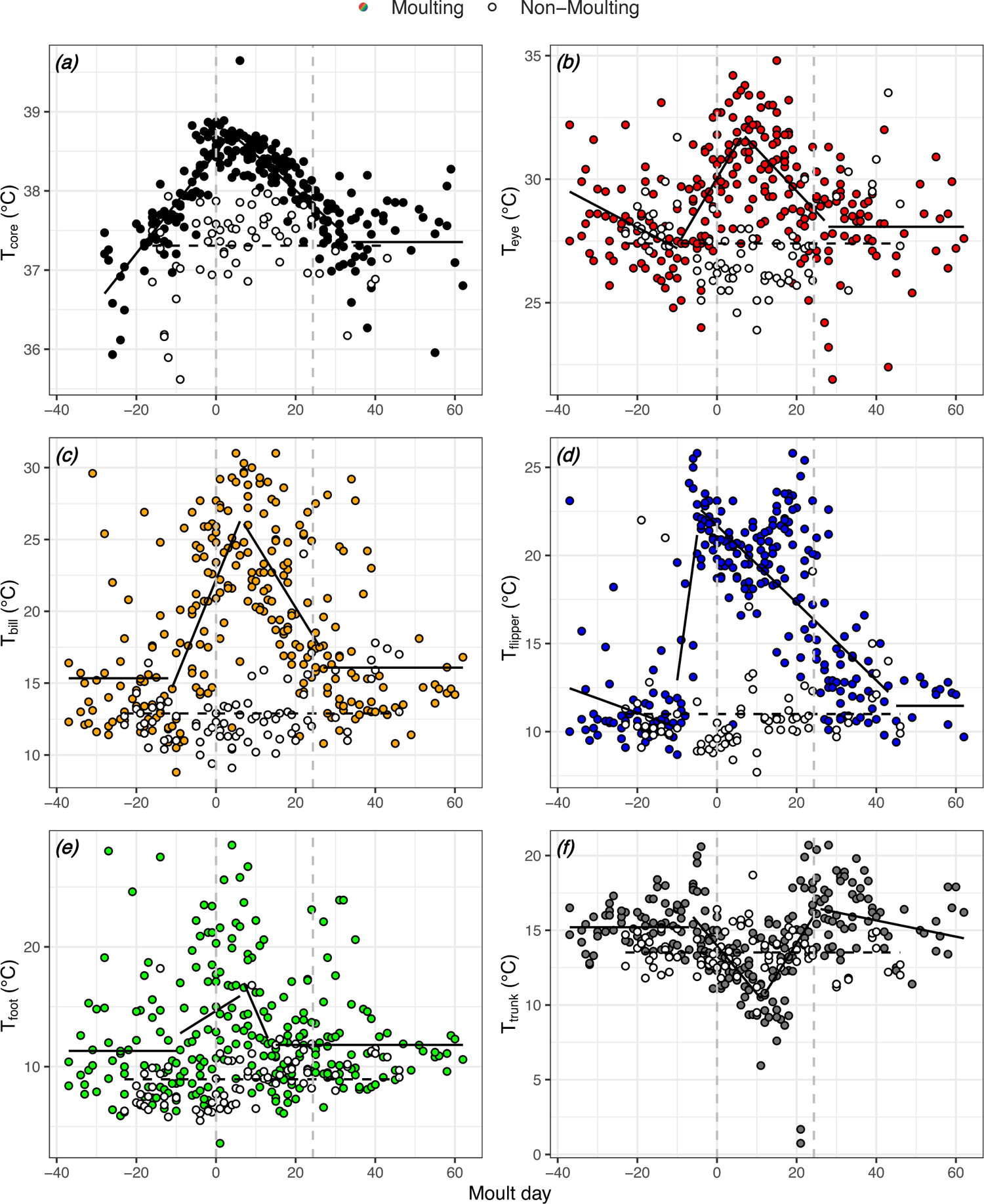
Variation in core (a) and surface temperatures (b: eye, c: bill, d: flipper, e: foot and f: trunk) of captive king penguins in relation to moult dynamics. The dashed line at day 0 indicates the first visible signs of moult, and the one at ca. day 24 indicates the completion of plumage renewal. Coloured circles and solid lines represent moulting birds, whereas white circles and dashed lines represent non-moulting birds. Lines are predictions from the top ranked models (Table S1). n = 6 moulting and n = 2 non-moulting individuals.

### Wild king penguins

Wild king penguins were studied in 2023-24 (as part of a long-term observatory) in the ‘Baie du Marin’ colony on Possession Island (40), Crozet Archipelago, in the Southern Indian Ocean (46°26’ S, 51°52’ E) from courtship in November 2023 to the 17^th^ of October 2024, when all scientific activities were shut down because of an outbreak of highly pathogenic avian influenza (HPAI) at the study site (41). At breeding initiation (Nov-Dec 2023), 48 breeding pairs were temporarily marked with hair dye on the breast feathers (*i.e.* lasting until the post-breeding moult) and permanently marked with thermosensitive pit-tags (BioTherm13 tag, resolution ± 0.1°C, accuracy ± 0.5°C, between 33°C and 43°C, Biomark, USA) in the inter-scapular region. These thermosensitive pit-tags enable long-term individual identification and subcutaneous temperature measurements (22). Subcutaneous temperature measurements were conducted during incubation (3^rd^ day of an incubation shift on land) as part of other long-term monitoring procedures, in less than 3 minutes after capture to avoid handling-stress induced changes in body temperature (40). A subsample of birds was equipped with core temperature ingestible loggers (Bodycap, Anipill, France, sampling rate = 5 or 15 min) during incubation. At the time of post-breeding moult (Sep-Oct 2024), the colony was searched daily to identify marked penguins being back on land to moult (N = 43 penguins observed 1 to 10 times during moult), scoring their moult according to ((10); M1 to M7 stages), noting the presence/absence of panting behaviour (22) and the usage of the river as a potentially cooling substrate (*i.e.* feet in the river: yes/no). Approximately on their 3^rd^ day on land, 3 moulting individuals (*i.e.* stage M1 for two individuals and stage M2 for another one) were recaptured, had their subcutaneous temperature recorded in less than 3 minutes after capture, and were equipped with a core temperature ingestible logger. At moult completion (stage M7), individuals were re-captured to download core body temperature data, which was unfortunately only possible for one individual before field site closure linked to HPAI outbreak. Considering the limited sample size, no statistical tests were performed and individual data points are presented. To quantify the environmental heat load faced by wild *vs.* captive individuals (*i.e.* including solar radiation and wind speed in addition to air temperature and humidity used for T_w_ in captivity), Wet Bulb Globe Temperature (WBGT) values were computed using the R package HeatStress function wbgt.Liljegren(). Environmental heat load in the wild was lower during moult than breeding, and captive penguins experienced an environmental heat load being intermediate between moulting and breeding individuals in the wild (Fig. S1).

## Results

### Captive penguins

Moulting and non-moulting birds were never observed panting during the study. The visible part of moult lasted on average 24.3 ± 0.9 days in captive individuals.

Changes in body temperatures during moult followed a non-linear pattern (Fig. 1). While T_core_ was best fitted with 2 breakpoints (*i.e.* an increase followed by a decrease during moult and then stabilisation post-moult, Fig 1a, Table S1), all surface temperatures were best fitted with at least 3 breakpoints that corresponded to 3 main phases: pre-moult, moult (with 2 segments surrounding peak-moult: early-moult and late-moult) and post-moult (Fig. 1b to 1f). While surface temperatures (T_eye_, T_bill_, T_flipper_, T_foot_ and T_trunk_) during pre- and post-moult periods were not always constant, the magnitude of changes during these periods were limited (Fig. 1 and Table S1). During moult, T_eye_, T_bill_, T_flipper_ and T_foot_ initially increased during early-moult (although non-significantly for T_foot_) and then decreased during late-moult, so post-moult levels were comparable to pre-moult and non-moulting levels (Fig. 1b to 1e; Fig 2b to 2e; Table S1). Conversely, T_trunk_ (*i.e.* an inverse proxy of insulation provided by the plumage) initially decreased and then increased back to levels comparable to pre-moult and non-moulting levels (Fig. 1f and 2f, Table S1).

**Figure 2.**
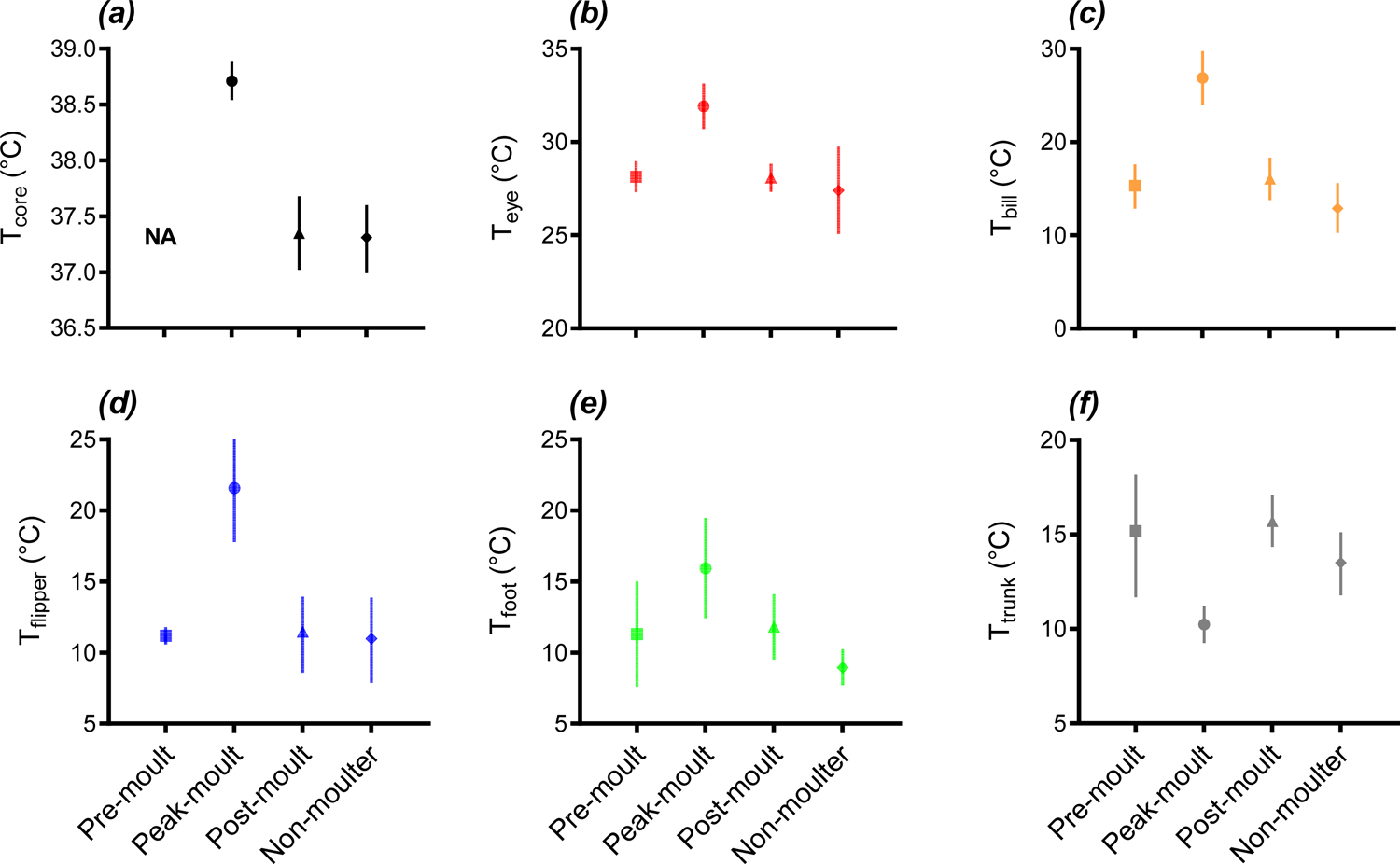
Predicted core (a) and surface body temperatures (b: eye, c: bill, d: flipper, e: foot and f: trunk) of captive king penguins at key stages during moult (i.e. pre-moult, peak-moult and post-moult based on breakpoint analyses) and in non-moulting individuals. Error bars represent 95% CI; n = 2-6 moulting and n = 2 non-moulting individuals.

Despite birds being equipped with a T_core_ logger on average 16.5 ± 4.8 days before visible changes in plumage occurred (*i.e.* moult day 0), we did not detect a pre-moulting stable period for T_core_ (Fig. 1a). Thus, we do not report a predicted T_core_ for the pre-moult stage (Fig. 2a), and we instead draw inferences by comparing T_core_ values between peak-moult and post-moult. T_core_ was *ca.* 1.3°C higher during peak moult than post-moult or than in non-moulting individuals. Body temperatures were overall mostly indistinguishable between pre-moult, post-moult and non-moulting individuals (Fig. 2). The changes in surface temperatures during moult were drastic, with an increase at peak moult of *ca.* +4.0°C for T_eye_ (Fig. 2b), +11.5°C for T_bill_ (Fig. 2c), +10.2°C for T_flipper_ (Fig. 2d), +5.0°C for T_foot_ although 95% CI overlapped with most other stages (Fig. 2e), and a decrease of *ca.* −5.0°C for T_trunk_ (Fig. 2f).

For body surface temperatures, the initial rise (*i.e.* moult start) in T_eye_, T_bill_, T_flipper_ and T_foot_ occurred simultaneously around 10 days before moult day 0 (Fig. 3). The initial drop in T_trunk_ occurred slightly later (*ca.* moult day −6), but not significantly so considering 95% CI overlapped (Fig. 3). The peak in T_flipper_ was reached approximately 10 days earlier than the peaks in T_core_, T_eye_, T_bill_, and T_foot_, while the peak in plumage insulation (*i.e.* lowest T_trunk_) occurred approximately 5 days later (Fig. 3, non-overlapping 95% CI). The peak in T_core_ occurred approximately 3 days before the peaks In T_eye_, T_bill_ and T_foot_, but not significantly so considering 95% CI overlapped (Fig. 3). T_foot_ was the first to return to baseline levels at *ca.* moult day 14, followed by T_trunk_, T_bill_, T_eye_ (*ca.* moult day 27), then T_core_ (*ca.* moult day 34) and then only by T_flipper_ (*ca.* moult day 44). While 95% CI overlapped between T_core_ and T_bill_, T_eye_, T_flipper_, this was not the case between T_core_ and T_trunk_ (Fig. 3).

**Figure 3.**
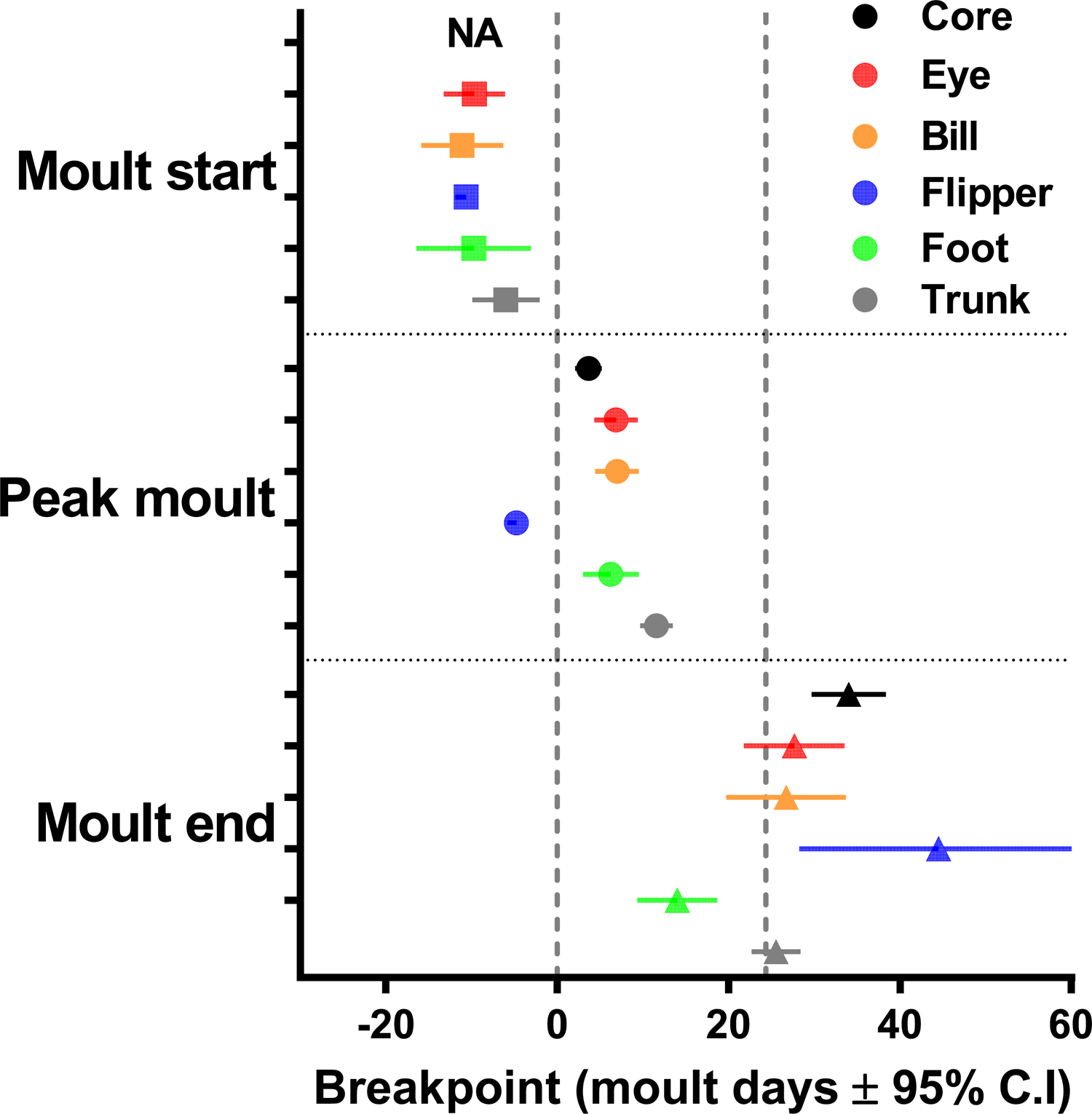
Temporal dynamics of changes in body temperatures during moult in captive king penguins based on breakpoint analyses. The dashed line at day 0 indicates the first visible signs of moult, and the one at ca. day 24 indicates the average time to visual completion of plumage renewal. Error bars represent 95% CI, N = 6 captive individuals.

### A glimpse at the natural scenario

Out of 87 observations from 43 wild individuals (mostly during early moulting stages), none were observed panting, whereas breeding individuals have been reported to pant during *ca.* 20% of behavioural observations in the same population (22). Moulting birds immersed their feet in the river during 75.8% of our observations. The visible part of moult lasted 13 days in the only individual that completed its moult before the HPAI-related closure of the field site.

T_subcut_ was approximately 1.4°C higher in early-moulting birds compared to the same individuals measured while breeding (Fig. 4a). Average daily T_core_ was overall 1.2°C higher during moulting compared to breeding in one wild individual (Fig. 4b), despite environmental heat load (*i.e.* WBGT) being lower during moult (Fig. 4c) and this individual being consistently observed with its feet immersed in the river. This wild individual had T_core_ within the range of captive non-moulting individuals during breeding (except the first day), and within the range of captive peak-moult between approximately 4 and 11 days of its moulting fast on land (Fig. 4b). T_core_ of this moulting individual did not return to breeding or captive non-moulting values, even once moult was visibly completed (Fig. 4b).

**Figure 4.**
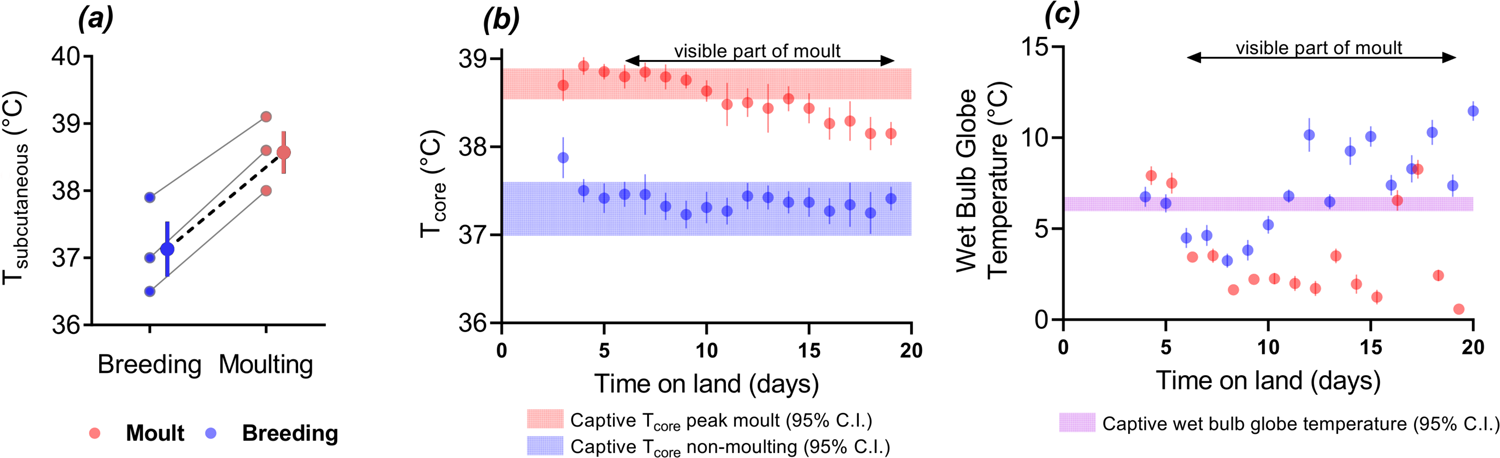
Subcutaneous (a) and core (b) body temperatures of moulting vs. breeding wild king penguin(s), along with (c) corresponding environmental heat load measured as Wet Bulb Globe Temperature. The same individual(s) were measured during incubation and moulting within the same breeding cycle (2023–24). (a) N = 3 individuals measured during incubation and early moult (ca. 3 day of fasting on land), with individual values presented in colour with grey lines, and mean ± SE in colour with black dashed line. (b) N = 1 individual measured during incubation (blue) and moulting (red) within the same breeding cycle (2023–24), circles and error bars represent daily average core body temperature ± SE, and the black arrow represents the visible part of moult (i.e. from new feathers emergence to fully new plumage). For comparison to results from captive individuals, the shaded areas represent the 95% CI of core body temperature for captive non-moulters (blue, n = 2) and during peak moult (red, n = 6). (c) Circles and error bars represent daily average Wet Bulb Globe Temperature ± SE during incubation (blue) and moulting (red), and the shaded purple area represent the 95% CI of Wet Bulb Globe Temperature for captive individuals for comparison.

## Discussion

### Warming up to a new coat

Our results provide evidence that moulting elicits thermovascular responses that increase heat loss in king penguins, as previously described in gentoo penguins (10). We now also provide evidence that core body temperature in king penguins markedly increases during moult (*i.e.* up to +1.4°C and +1.2°C in captive and wild birds respectively), suggesting either a failure to dissipate excess heat generated by the up-regulation of metabolism (*thermal challenge hypothesis*), or a regulated increase of body temperature during moult (*warm-up hypothesis*). By assessing the timing of changes in surface and core body temperature, we found that surface temperatures increased rapidly, and likely after the rise in core temperature occurred, thereby providing support for the *warm-up* hypothesis. Coupled with the lack of panting observed in the study period, these findings suggest that increasing peripheral heat dissipation was likely enough to maintain core temperature below a maximum threshold. The limited evidence we gathered from wild king penguins (*i.e.* +1.4°C in subcutaneous and +1.2°C in core body temperature) suggests that the hyperthermic pattern we observed in captivity also occurs under natural conditions, where behavioural heat dissipation was less constrained because individuals had access to cold water, and used it for partial immersion.

King penguins must accomplish moulting efficiently to avoid reproductive failure because food availability decreases rapidly after the moulting period (42). It is known that king penguins that arrive from the sea later in the season initiate feather loss more quickly and undergo a contracted moult period relative to individuals that arrive earlier (43). Hyperthermia may benefit moulting penguins by accelerating the process of feather growth, thus enabling them to better match high food availability. Assuming that king penguins in our study had a Q10 equal to 2.3 (Roussel et al., *pers. Com.*; 44) the observed increase in body temperature of ∼1.4°C would yield a 12.4% increase in metabolic rate, which may shorten moult by *ca.* 2.5 days, equal to a 9.3% reduction in the total duration of moult.

Positive correlations between feather growth and body temperature have been observed in chickens, where individuals with an early feathering genotype display higher body temperatures and faster feather growth during the first six weeks post-hatching (45). A similar pattern has been observed indirectly in turkeys, where individuals held at higher ambient temperature had longer feathers at 14 days post-hatching (46), although this may partially be explained by metabolic costs being diverted to thermoregulation instead of feather growth at the lower temperature treatment used in that study. Further supporting the association between high body temperature and accelerated feather growth, the majority of feather growth occurs during the day when body temperature and metabolic rate are highest (47). In penguins, moulting-induced hyperthermia occurs alongside increases in blood perfusion at the surface ((10); this study), which likely helps to deliver nutrients to peripheral tissues that is necessary for feather growth (48).

### Thermoregulation in captivity and in the wild during catastrophic moult

Core and surface temperatures in captive non-moulting birds remained stable during the study period relative to moulting birds, which exhibited distinct changes over time. Core temperature was the first to increase under captive conditions as we were not able to detect a pre-moulting stable period. During the gradual rise in core temperature, we observed increases in surface temperature in thermal windows (flippers, bill, and feet), indicating increased effort to dissipate heat. Interestingly, the flippers were the only thermal window to reach the maximum surface temperature before core body temperature reached its maximum, suggesting that the flippers may function as a first line of defense for peripheral heat dissipation in penguins, likely due to their large surface area and vascularization (49). On the other hand, the surface temperatures of the bill and feet closely tracked fluctuations in core temperature, as indicated by the similarities in the timing and direction of changes in these temperatures. That heat dissipation via thermal windows increased after core temperature had already been rising indicates that hyperthermia was likely regulated and maintained below a maximum threshold. This interpretation is further supported by the lack of panting observed throughout the study period. We found that the surface temperature of the trunk declined in captive moulting birds, indicating increased insulation due to the overlap of new and old layers of plumage (10, 50).

Despite the increased insulation during the moult, core temperature had already begun declining by the time insulation reached its maximum (i.e., the minimum of T_trunk_). This observation suggests that the rise in core temperature was primarily driven by increased metabolic heat generation, not by increased insulation as hypothesized in (10).

Conducting this research in captivity enabled us to eliminate variation that would be introduced by more dynamic thermal environments in the wild, thus isolating the effects of moulting on thermoregulation. On one hand, captivity prevented heating from solar radiation; on the other hand, captivity prevented cooling from wind chill and precipitation, and it limited behavioural thermoregulation in the form of habitat selection (*i.e.,* partial immersion in water). However, behavioural thermoregulation and environmental conditions, such as solar radiation, wind, and humidity remain essential considerations for wild animal thermoregulation (22, 51–53). Therefore, to provide insights about the relevance of our finding in captivity, we measured thermoregulatory behaviour, subcutaneous temperature, and core temperature in a limited number of wild king penguins. During 75.8% of our observations, free-living moulting king penguins had their feet immersed in the river, confirming that wild birds likely take advantage of additional means to dissipate heat. Despite this behaviour and greater potential for wind chill, core temperature measured in one wild moulting king penguin showed a similar hyperthermic pattern (+1.2°C compared to incubation) as the one found in captive moulting king penguins (+1.4°C). Although this result was obtained in only one individual, it is supported by the similar pattern observed in the subcutaneous temperature of three wild king penguins measured during both incubation and early-moult (+1.4°C). It is interesting to note that the core body temperature of captive individuals was similar to the wild scenario, both during (T_core_ ≃ 38.7°C) and outside (T_core_ ≃ 37.3°C) the moulting period, thereby suggesting that captivity does not markedly affect thermoregulation in king penguins.

### Seeking baselines: Operating points of body and surface temperatures

The data we report conform well to a negative feedback-based model of thermoregulation with multiple autonomous thermo-effectors (54). The elevation of metabolic heat generation while moulting represents a positive (i.e., heat) thermal challenge that ultimately triggers an increase in heat dissipation via the periphery. The rises in core, and subsequently surface, temperatures represent the changes in multiple operating points that would be necessary to balance heat loss with elevated heat generation. The thermal challenge introduced by moulting was large enough to elicit changes in peripheral heat loss, but not panting, suggesting that the thermoregulatory response observed lies within an inter-threshold zone (55–58). We interpret the difference in timing and magnitude of change in flipper temperature relative to other thermal windows to represent varying thresholds for response across body regions, a pattern supported by the specialization of regional heterothermy in birds (59), and indeed by the widespread presence of regional heterothermy in endotherms (60).

Previous studies have suggested that moulting poses a negative (i.e., cold) thermoregulatory challenge due to reduced insulation during feather loss, which is compensated by an increase in heat production (17–19, 61). If moulting poses a negative thermal challenge, one would expect a reduction in body temperature followed by upregulation of metabolic heat generation until the thermal challenge passes (i.e., hypothermia, not hyperthermia (54)). However, moulting in penguins [(62, 63), present study] and other birds (5, 64) can elicit elevations in body temperature (i.e., hyperthermia), the opposite of the pattern predicted if moulting was a negative thermal challenge. Moulting-induced hyperthermia may be less pronounced in species where heat loss occurs too efficiently for body temperature to be affected by upregulated metabolism (e.g., small passerines; (65)), or if the sampling period does not align with the onset of hyperthermia. In the wild, penguins arrive onshore with new feathers developed to between 18% to 42% of their final length (18, 66), and it is possible that studies in wild individuals may not have been able to detect early changes in core temperature occurring during the pre-moult or early-moult stages. Pre-moult elevation of core temperature is not mutually exclusive with increased energy cost of thermoregulation in later stages of the moult when insulation decreases quickly, as observed in (19).

Conducting this research in captivity enabled us to measure moulting and non-moulting individuals simultaneously throughout the entire moulting process. Despite starting to measure core temperature more than two weeks before the first visible change in plumage, our sampling period was too late to capture a stable pre-moult phase of core temperature. In gentoo penguins, lower surface temperatures were observed at the end of the study, possibly because the sampling period may have begun after body temperatures had already begun increasing (10). This and previous studies highlight the difficulty of establishing a baseline for normothermia, and these differences in sampling period may explain some of the differences between studies. We advise care when planning sampling periods for physiological studies of moulting because physiological changes in preparation for moulting that underly visible change in plumage may have an earlier start than what was previously appreciated in the literature.

We also advise care external and internal factors when measuring thermoregulatory traits, such as core temperature, surface temperatures, and thermoregulatory behaviors, because these can be influenced by multiple. For instance, acute stress can cause temperature changes in regions throughout the body (29, 33, 40, 67) and body temperatures also exhibit circadian patterns (68). Acute stress and circadian changes in body temperature could affect behaviours that vary with heat load, such as panting, a common thermoregulatory behavior in birds, including penguins (69, 70). In our system, we identified the daily visits of keepers as potential sources of error because of variation in timing, tasks, and levels of disturbance (underwater cleaning being less stressful than above-water cleaning). To avoid both stress-related and circadian influences on body temperatures and thermoregulation, we collected data at the same time each morning before keepers entered the enclosure. Yet, such potential issues are way more difficult to deal with in wild settings, where timing of measurement is not necessarily chosen by the experimenter, and acute stressors (*e.g.* aggressive interaction with neighbours, presence of predator) cannot be controlled and may not be noticed by the experimenter.

### Warming up in captivity – heating up in the wild?

We found that the increase in core temperature in moulting king penguins was likely regulated, as evidenced by the lack of influence from plumage insulation, the absence of panting behavior, and the delay in vascular perfusion. These patterns suggest that hyperthermia was likely employed as a regulated thermoregulatory strategy (71). Air temperature remained stable and far below body temperature during our study, and the warming effect of solar radiation was absent in captivity. Since environmental conditions are unlikely to be as forgiving in the wild, the current study raises concerns regarding the vulnerability of moulting penguins in the context of global climate change. High mortality rates have already been reported in multiple penguin species immediately after moulting due to the energetic challenge of this fasting period (18, 72–74). More recently, in 2019, a mortality event was recorded during early moult in little penguins following a heatwave on land (*Eudyptula minor*; (75)). Based on our finding that moulting poses a positive (i.e., heat) thermal challenge, we caution that moulting may increase the susceptibility of wild penguins to heat stress, especially as regions near the poles warm at a disproportionately rapid rate (24). Further investigation of thermoregulatory responses of penguins during moulting in the wild is thus deeply needed.

## Data availability and Electronic Supplementary Materials

The datasets, R code, and Electronic Supplementary Materials (Supplemental Figs. S0 and S1; Supplemental Tables S0-A, S0-B, and Table S1) used in this manuscript are available on FigShare: https://figshare.com/articles/dataset/Data_from_Warming_up_to_a_new_coat_moul_ng_king_penguins_exhibit_hyperthermia_and_increased_peripheral_heat_loss_/28629305

## Supporting information

Electronic Supplemental Materials

## Acknowledgments

We are grateful to Oceanopolis© and Dominique Barthelemy for their logistical support during the experiment. This work would not have been possible without the help of the animal care team: Christine Dumas, Alexiane Corcuff, Maxence Leroy, Mélanie Robert and Agathe Lefranc. We thank the members of polar program #119 (ECONERGY) JP Robin, VA Viblanc, P Bize, as well as to the French Polar Institute (IPEV) and the Terres Australes et Antarctiques Françaises for providing financial and logistical support for the study of wild king penguins. We are grateful to Marine Montblanc, Norith Eckbo, Samuel Laporte, Zohria-Lys Guillerm, Camille De Pasquale and the entire Alfred Faure field station for their help in the field. We thank Raymond M. Danner for providing financial and logistical support to JDZ, and for his helpful comments on the proposal. We thank the following students for their assistance in extracting data from thermal images: Abby E. Lehman, Olivia P. Wanex, Roberto Bonifacio-Dominguez, Tyler W. Vanwinkle. We thank Glenn J. Tattersall and Dominic J. McCafferty for providing feedback regarding standardization of the thermal camera measurements, as well as D. Roussel for providing unpublished Q10 values of king penguins. The wild part of this study was part of the long-term Studies in Ecology and Evolution (SEE-Life) program of the CNRS. AL was supported by ISblue project, Interdisciplinary graduate school for the blue planet (ANR-17-EURE-0015) and co-funded by a grant from the French government under the program “Investissements d’Avenir” embedded in France 2030. This project received financial support from the “Région Bretagne”. The project benefitted from an IdEx grant from the Université de Strasbourg (HotPenguin) granted to AS. EM was funded by the French Polar Institute. We thank the Company of Biologists and the Journal of Experimental Biology for providing a Traveling Fellowship to JDZ.

## Author’s contributions

Study design: ALew, AS, JZ. Funding acquisition: ALew, AS, ALey, JZ. Data collection in captivity: JZ, EP, ALew. Data collection in the field: EM, AS. Data analysis: JZ, EP, AS. Writing original draft: JZ, ALew and AS. Writing review and editing: ALey, EP, EM.

## Institutional Review Board Statement (IRB)

The study in captive penguins did not include invasive procedures. Scientific procedures for wild king penguins were approved by the French Ethical Committee (APAFIS#31268-2021042117037897 v3) and the Terres Australes et Antarctiques Françaises (Arrêtés TAAF A-2023-88).

## New and Noteworthy

Penguins may experience heat stress while moulting, which causes increased metabolic heat generation and insulation. We assessed thermoregulation in moulting captive king penguins (Aptenodytes patagonicus) using simultaneous measurements of core and surface temperatures. By measuring temperature throughout the entirety of the moult, we found that hyperthermia and increased peripheral heat dissipation are prevalent in moulting king penguins. We caution that moulting-induced hyperthermia may contribute to the susceptibility of penguins to heat stress in the wild.

## References

1. Beltran RS, Burns JM, Breed GA. Convergence of biannual moulting strategies across birds and mammals. Proc R Soc B 285: 20180318, 2018. doi: 10.1098/rspb.2018.0318.

2. Terrill RS, Shultz AJ. Feather function and the evolution of birds. Biological Reviews 98: 540–566, 2023. doi: 10.1111/brv.12918.

3. Jenni L, Winkler R. The Biology of Moult in Birds. London: HELM, 2020.

4. Carey C, editor. Energetics and Nutrition of Molt. In: Avian Energetics and Nutritional Ecology. Springer US, p. 158–198.

5. Lustick S. Energy Requirements of Molt in Cowbirds. The Auk 87: 742–746, 1970. doi: 10.2307/4083708.

6. Dietz MW, Daan S, Masman D. Energy Requirements for Molt in the Kestrel Falco tinnunculus. Physiological Zoology 65: 1217–1235, 1992. doi: 10.1086/physzool.65.6.30158276.

7. Wijnandts H. Ecological Energetics of the Long-Eared Owl (Asio Otus). Ardea 38–90: 1– 92, 2002. doi: 10.5253/arde.v72.p1.

8. Le Maho Y, Delclitte P, Chatonnet J. Thermoregulation in fasting emperor penguins under natural conditions. American Journal of Physiology-Legacy Content 231: 913–922, 1976. doi: 10.1152/ajplegacy.1976.231.3.913.

9. Gavrilov V. Metabolism of molting birds. Zoologicheskii Zhurnal 53: 1363–1375, 1974.

10. Lewden A, Halna Du Fretay T, Stier A. Changes in body surface temperature reveal the thermal challenge associated with catastrophic moult in captive gentoo penguins. Journal of Experimental Biology 227: jeb247332, 2024. doi: 10.1242/jeb.247332.

11. Kazas S, Benelly M, Golan S. The Humboldt Penguin (Spheniscus humboldti) Rete Tibiotarsale – A supreme biological heat exchanger. Journal of Thermal Biology 67: 67–78, 2017. doi: 10.1016/j.jtherbio.2017.04.011.

12. Frost PGH, Siegfried WR, Greenwood PJ. Arterio-venous heat exchange systems in the Jackass penguin Spheniscus demersus. Journal of Zoology 175: 231–241, 1975. doi: 10.1111/j.1469-7998.1975.tb01398.x.

13. Thomas DB, Fordyce RE. Biological Plasticity in Penguin Heat-Retention Structures. The Anatomical Record 295: 249–256, 2012. doi: 10.1002/ar.21538.

14. Enstipp MR, Bost C-A, Le Bohec C, Bost C, Le Maho Y, Weimerskirch H, Handrich Y. Apparent changes in body insulation of juvenile king penguins suggest an energetic challenge during their early life at sea. Journal of Experimental Biology 220: 2666–2678, 2017. doi: 10.1242/jeb.160143.

15. Ainley DG, Wilson RP. Hot Penguins: Cold Water. In: The Aquatic World of Penguins. Springer International Publishing, p. 217–256.

16. De Vries J, Van Eerden MR. Thermal Conductance in Aquatic Birds in Relation to the Degree of Water Contact, Body Mass, and Body Fat: Energetic Implications of Living in a Strong Cooling Environment. Physiological Zoology 68: 1143–1163, 1995. doi: 10.1086/physzool.68.6.30163797.

17. Adams NJ, Brown CR. Energetics of Molt in Penguins. In: Penguin Biology. Elsevier, p. 297–315.

18. Cherel Y, Charrassin JB, Challet E. Energy and protein requirements for molt in the king penguin Aptenodytes patagonicus. American Journal of Physiology-Regulatory, Integrative and Comparative Physiology 266: R1182–R1188, 1994. doi: 10.1152/ajpregu.1994.266.4.R1182.

19. Baudinette R, Gill P, O’driscoll M. Energetics of the Little Penguin, Eudyptula Minor: Temperature Regulation, the Calorigenic Effect of Food, and Moulting. Aust J Zool 34: 35, 1986. doi: 10.1071/ZO9860035.

20. Groscolas R, Cherel Y. How to Molt While Fasting in the Cold: The Metabolic and Hormonal Adaptations of Emperor and King Penguins. Ornis Scandinavica 23: 328, 1992. doi: 10.2307/3676657.

21. Stonehouse B. The General Biology and Thermal Balances of Penguins. In: Advances in Ecological Research. Elsevier, p. 131–196.

22. Noiret A, Lewden A, Lemonnier C, Bocquet C, Montblanc M, Bertile F, Hoareau M, Marcon E, Robin J-P, Viblanc VA, Stier A. Mind the polar sun: Solar radiations trigger frequent heat stress in breeding king penguins, despite relatively cool air temperatures. Evolutionary Biology: 2024.

23. Welman S, Pichegru L. Nest microclimate and heat stress in African Penguins Spheniscus demersus breeding on Bird Island, South Africa. Bird Conservation International 33: e34, 2023. doi: 10.1017/S0959270922000351.

24. Nel W, Hedding DW, Rudolph EM. The sub-Antarctic islands are increasingly warming in the 21st century. Antarctic Science 35: 124–126, 2023. doi: 10.1017/S0954102023000056.

25. Gorodetskaya IV, Durán-Alarcón C, González-Herrero S, Clem KR, Zou X, Rowe P, Rodriguez Imazio P, Campos D, Leroy-Dos Santos C, Dutrievoz N, Wille JD, Chyhareva A, Favier V, Blanchet J, Pohl B, Cordero RR, Park S-J, Colwell S, Lazzara MA, Carrasco J, Gulisano AM, Krakovska S, Ralph FM, Dethinne T, Picard G. Record-high Antarctic Peninsula temperatures and surface melt in February 2022: a compound event with an intense atmospheric river. npj Clim Atmos Sci 6: 202, 2023. doi: 10.1038/s41612-023-00529-6.

26. Davis LS, Darby JT. Penguin biology. San Diego New York Boston [etc.]: Academic press, 1990.

27. Froget G, Handrich Y, Maho YL, Rouanet J-L, Woakes AJ, Butler PJ. The heart rate/oxygen consumption relationship during cold exposure of the king penguin: a comparison with that during exercise. Journal of Experimental Biology 205: 2511–2517, 2002. doi: 10.1242/jeb.205.16.2511.

28. Cherel Y, Charrassin J-B, Handrich Y. Comparison of Body Reserve Buildup in Prefasting Chicks and Adults of King Penguins (Aptenodytes patagonicus). Physiological Zoology 66: 750–770, 1993. doi: 10.1086/physzool.66.5.30163822.

29. Tabh JKR, Burness G, Wearing OH, Tattersall GJ, Mastromonaco GF. Infrared thermography as a technique to measure physiological stress in birds: Body region and image angle matter. Physiol Rep 9, 2021. doi: 10.14814/phy2.14865.

30. Jerem P, Jenni-Eiermann S, Herborn K, McKeegan D, McCafferty DJ, Nager RG. Eye region surface temperature reflects both energy reserves and circulating glucocorticoids in a wild bird. Sci Rep 8: 1907, 2018. doi: 10.1038/s41598-018-20240-4.

31. R Core Team. R: A Language and Environment for Statistical Computing [Online]. R Foundation for Statistical Computing: 2025. https://www.R-project.org/.

32. Casanueva A. HeatStress. Zenodo: 2019.

33. Zuluaga JD, Danner RM. Acute stress and restricted diet reduce bill-mediated heat dissipation in the song sparrow (Melospiza melodia): implications for optimal thermoregulation. Journal of Experimental Biology 226: jeb245316, 2023. doi: 10.1242/jeb.245316.

34. Muggeo VMR. segmented: Regression Models with Break-Points / Change-Points Estimation (with Possibly Random Effects). 2.1-4, 2003.

35. Bates D, Mächler M, Bolker B, Walker S. Fitting Linear Mixed-Effects Models Using lme4. J Stat Soft 67, 2015. doi: 10.18637/jss.v067.i01.

36. Burnham KP, Anderson DR, editors. Model Selection and Multimodel Inference. Springer New York.

37. Bartoń K. MuMIn: Multi-Model Inference. 1.48.11, 2010.

38. Lüdecke D. ggeffects: Tidy Data Frames of Marginal Effects from Regression Models. JOSS 3: 772, 2018. doi: 10.21105/joss.00772.

39. Wickham H. ggplot2: Elegant Graphics for Data Analysis. Springer New York.

40. Lewden A, Ward C, Noiret A, Avril S, Abolivier L, Gérard C, Hammer TL, Raymond É, Robin J-P, Viblanc VA, Bize P, Stier A. Surface temperatures are influenced by handling stress independently of corticosterone levels in wild king penguins (Aptenodytes patagonicus). Journal of Thermal Biology 121: 103850, 2024. doi: 10.1016/j.jtherbio.2024.103850.

41. Clessin A, Briand F-X, Tornos J, Lejeune M, Pasquale CD, Fischer R, Souchaud F, Hirchaud E, Bralet T, Guinet C, McMahon CR, Grasland B, Baele G, Boulinier T. Mass mortality events in the sub-Antarctic Indian Ocean caused by long-distance circumpolar spread of highly pathogenic avian influenza H5N1 clade 2.3.4.4b. Microbiology: 2025.

42. Olsson O. Seasonal Effects of Timing and Reproduction in the King Penguin: A Unique Breeding Cycle. Journal of Avian Biology 27: 7, 1996. doi: 10.2307/3676955.

43. Gauthier-Clerc M, Le Maho Y, Gendner J-P, Handrich Y. Moulting fast and time constraint for reproduction in the king penguin. Polar Biol 25: 288–295, 2002. doi: 10.1007/s00300-001-0342-y.

44. Monternier P-A, Marmillot V, Rouanet J-L, Roussel D. Mitochondrial phenotypic flexibility enhances energy savings during winter fast in king penguin chicks. J Exp Biol 217: 2691–2697, 2014. doi: doi:10.1242/jeb.104505.

45. Noubandiguim M, Erensoy K, Sarica M. Feather growth, bodyweight and body temperature in broiler lines with different feathering rates. SA J An Sci 51, 2021. doi: 10.4314/sajas.v51i1.10.

46. Wylie LM, Robertson GW, MacLeod MG, Hocking PM. Effects of ambient temperature and restricted feeding on the growth of feathers in growing turkeys. British Poultry Science 42: 449–455, 2001. doi: 10.1080/00071660120070631.

47. Lillie FR, Wang H. Physiology of Development of the Feather: IV. The Diurnal Curve of Growth in Brown Leghorn Fowl. Proc Natl Acad Sci USA 26: 67–85, 1940. doi: 10.1073/pnas.26.1.67.

48. Lillie FR. Physiology of Development of the Feather. III Growth of the Mesodermal Constituents and Blood Circulation in the Pulp. Physiological Zoology 13: 143–176, 1940. doi: 10.1086/physzool.13.2.30151536.

49. Lewden A, Nord A, Bonnet B, Chauvet F, Ancel A, McCafferty DJ. Body surface rewarming in fully and partially hypothermic king penguins. J Comp Physiol B 190: 597– 609, 2020. doi: 10.1007/s00360-020-01294-1.

50. Le Maho Y. The Emperor Penguin: A Strategy to Live and Breed in the Cold: Morphology, physiology, ecology, and behavior distinguish the polar emperor penguin from other penguin species, particularly from its close relative, the king penguin. American Scientist 65: 680–693, 1977.

51. Freeman MT, Coulson B, Short JC, Ngcamphalala CA, Makola MO, McKechnie AE. Evolution of avian heat tolerance: The role of atmospheric humidity. Ecology 105: e4279, 2024. doi: 10.1002/ecy.4279.

52. Wolf BO, Walsberg GE. Thermal Effects of Radiation and Wind on a Small Bird and Implications for Microsite Selection. Ecology 77: 2228–2236, 1996. doi: 10.2307/2265716.

53. Cunningham SJ, Gardner JL, Martin RO. Opportunity costs and the response of birds and mammals to climate warming. Frontiers in Ecology and the Environment 19: 300–307, 2021.

54. Mitchell D, Fuller A, Snelling EP, Tattersall GJ, Hetem RS, Maloney SK. Revisiting concepts of thermal physiology: understanding negative feedback and set-point in mammals, birds, and lizards. Biological Reviews 100: 1317–1346, 2025. doi: 10.1111/brv.70002.

55. Bligh J. The thermosensitivity of the hypothalamus and thermoregulation in mammals. Biological Reviews 41: 317–365, 1966. doi: 10.1111/j.1469-185X.1966.tb01496.x.

56. Notley SR, Mitchell D, Taylor NAS. A century of exercise physiology: concepts that ignited the study of human thermoregulation. Part 3: Heat and cold tolerance during exercise. Eur J Appl Physiol 124: 1–145, 2024. doi: 10.1007/s00421-023-05276-3.

57. Taylor NAS, Gordon CJ. The origin, significance and plasticity of the thermoeffector thresholds: Extrapolation between humans and laboratory rodents. Journal of Thermal Biology 85: 102397, 2019. doi: 10.1016/j.jtherbio.2019.08.003.

58. Mitchell D, Snelling EP, Hetem RS, Maloney SK, Strauss WM, Fuller A. Revisiting concepts of thermal physiology: Predicting responses of mammals to climate change. Journal of Animal Ecology 87: 956–973, 2018. doi: 10.1111/1365-2656.12818.

59. McQueen A, Barnaby R, Symonds MRE, Tattersall GJ. Birds are better at regulating heat loss through their legs than their bills: implications for body shape evolution in response to climate. Biol Lett 19: 20230373, 2023. doi: 10.1098/rsbl.2023.0373.

60. Angilletta M, Cooper B, Schuler M, Boyles J. The evolution of thermal physiology in endotherms. Front Biosci E2: 861–881, 2010. doi: 10.2741/e148.

61. Green JA, Butler PJ, Woakes AJ, Boyd IL. Energetics of the moult fast in female macaroni penguins Eudyptes chrysolophus. Journal of Avian Biology 35: 153–161, 2004. doi: 10.1111/j.0908-8857.2004.03138.x.

62. Farner DS. Incubation and Body Temperatures in the Yellow-eyed Penguin. The Auk 75: 249–262, 1958.

63. Groscolas R. Study of molt pasting followed by an experimental forced fasting in the emperor penguin Aptenodytes forsteri: Relationship between feather growth, body weight loss, body temperature and plasma fuel levels. Comparative Biochemistry and Physiology Part A: Physiology 61: 287–295, 1978. doi: 10.1016/0300-9629(78)90111-1.

64. Newton I. The Temperatures, Weights, and Body Composition of Molting Bullfinches. The Condor 70: 323–332, 1968. doi: 10.2307/1365926.

65. Chilgren J. Dynamics and bioenergetics of postnuptial molt in captive White-crowned Sparrows (Zonotrichia leucophrys gambelii). Washington State University: 1975.

66. Brown CR. Feather growth, mass loss and duration of moult in macaroni and rock-hopper penguins. Ostrich 57: 180–184, 1986. doi: 10.1080/00306525.1986.9633647.

67. Herborn KA, Jerem P, Nager RG, McKeegan DEF, McCafferty DJ. Surface temperature elevated by chronic and intermittent stress. Physiology & Behavior 191: 47–55, 2018. doi: 10.1016/j.physbeh.2018.04.004.

68. Prinzinger R, Preßmar A, Schleucher E. Body temperature in birds. Comparative Biochemistry and Physiology Part A: Physiology 99: 499–506, 1991. doi: 10.1016/0300-9629(91)90122-S.

69. Murrish DE. Acid-Base Balance in Three Species of Antarctic Penguins Exposed to Thermal Stress. Physiological Zoology 55: 137–143, 1982.

70. Welman S, Green JA, Ryan PG, Parsons NJ, Pichegru L. Body temperature and thermoregulatory behaviour in the Endangered African Penguin Spheniscus demersus. Bird Conservation International 34: e29, 2024. doi: 10.1017/S095927092400025X.

71. Gerson AR, McKechnie AE, Smit B, Whitfield MC, Smith EK, Talbot WA, McWhorter TJ, Wolf BO. The functional significance of facultative hyperthermia varies with body size and phylogeny in birds. Functional Ecology 33: 597–607, 2019. doi: 10.1111/1365-2435.13274.

72. Heezik YV, Davis L. Effects of food variability on growth rates, fledging sizes and reproductive success in the Yellow-eyed Penguin Megadyptes antipodes. Ibis 132: 354– 365, 1990. doi: 10.1111/j.1474-919X.1990.tb01055.x.

73. Boersma PD. Breeding Patterns of Galápagos Penguins as an Indicator of Oceanographic Conditions. Science 200: 1481–1483, 1978. doi: 10.1126/science.200.4349.1481.

74. Dann P, Cullen JM, Thoday R, Jessop R. Movements and Patterns of Mortality at Sea of Little Penguins Eudyptula minor from Phillip Island, Victoria. Emu - Austral Ornithology 91: 278–286, 1991. doi: 10.1071/MU9910278.

75. Tworkowski L, Dann P, Ellenburg U, Robert K. Impacts of terrestrial heat waves on survival of little penguins during moult. 2021.

