## Supplementary material for "Warming up to a new coat: moulting king penguins exhibit hyperthermia and increased peripheral heat loss": Electronic Supplemental Materials

**Electronic Supplementary Material (ESM)**

**Supplementary methods S0: Standardization between IR cameras**

We standardized measurements made with the three cameras using two separate linear models that were built using the surface temperature of the plumage of non-moulting birds. We selected the plumage because of its lack of vascularization to avoid confounding differences between cameras with natural variation in peripheral vasoconstriction, and we used data from non-moulting birds to avoid confounding differences between cameras with changes associated with the moulting period. The linear models included the camera as a fixed effect to quantify the effect of each camera on the measured surface temperature, and T_w_ to account for variation introduced from environmental conditions (temperature and humidity). We built the first regression using data collected daily during the study when one camera was available at a time (*i.e.*, non-simultaneous comparisons), and we built the second regression using data collected at the end of the experiment when all three cameras were available for simultaneous comparisons. Both models yielded similar results, and we made corrections using the parameter estimates from the models built with simultaneous comparisons because we expect that this dataset is more resistant to temporal variation in surface temperature.

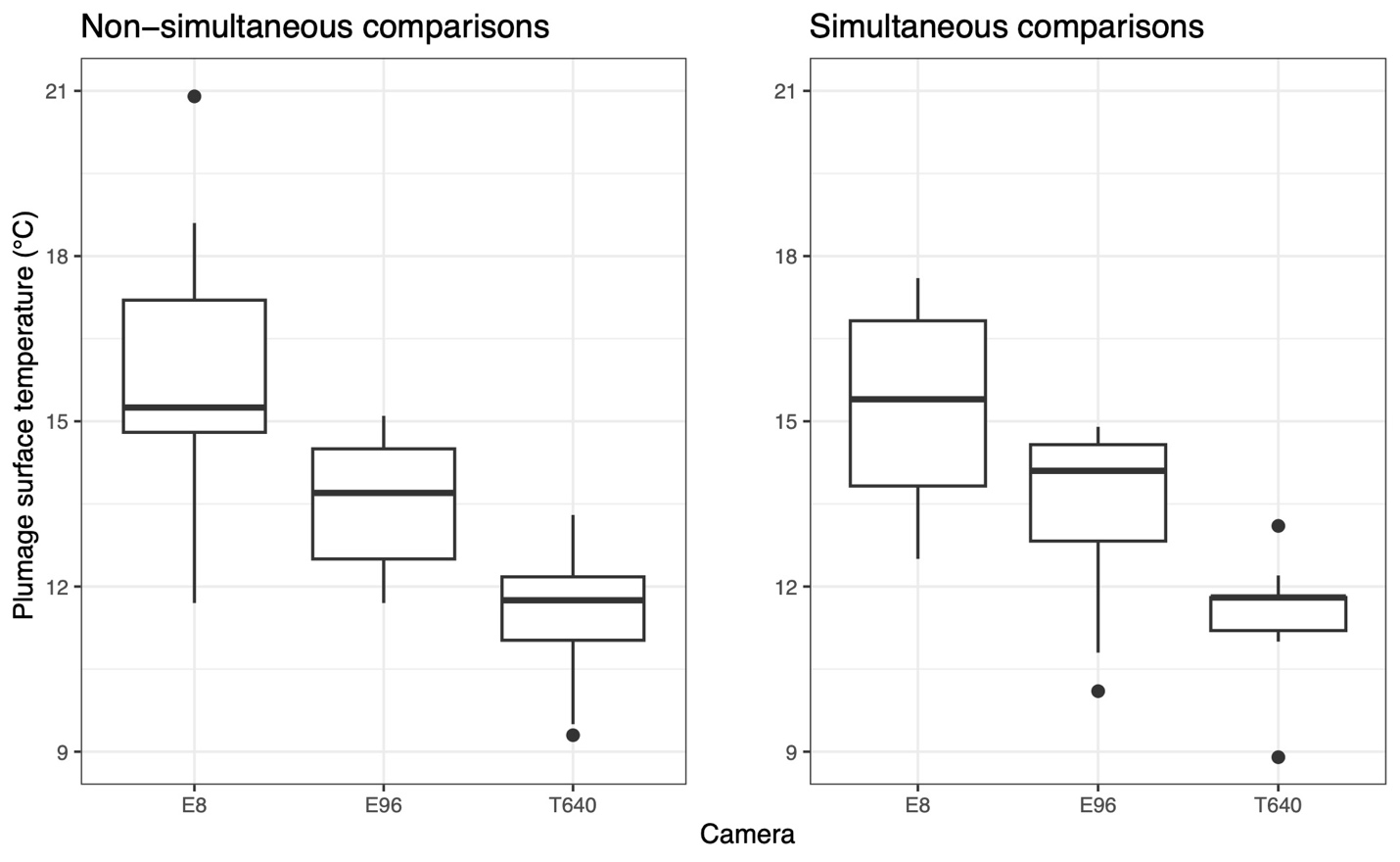

**Figure S0. Differences in surface temperature measurements between cameras.** Data are of plumage temperature over the course of the study (left) and at the end of the study when all three cameras were available for simultaneous comparisons (right).

**Table S0-A.** Parameter estimates for the effects of the two replacement cameras on surface temperature measurements. Linear models were built using simultaneous comparisons between the E96 and each of the replacements (E8 and T640), and the resulting parameter estimates were used to standardize measurements made with the different cameras.

| **Camera** | **Parameter estimate** | **Standard error** | **p-value** |
| --- | --- | --- | --- |
| E8 | 2.20 | 0.594 | 0.0138 |
| T640 | -1.90 | 0.389 | 0.0164 |

**Table S0-B.** Resolutions of the three thermal imaging cameras used in this study.

| **Camera** | **Resolution** |
| --- | --- |
| E96 | 640X480 |
| E8 | 320X240 |
| T640 | 640X480 |

**Supplementary results S1:**

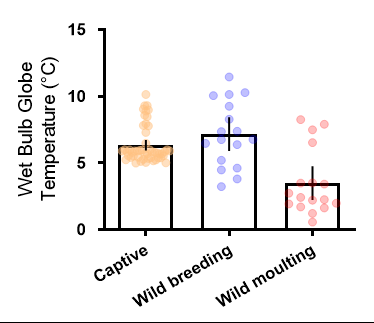

**Figure S1.** **Environmental heat load evaluated as Wet Bulb Globe Temperature (WBGT) between captive conditions (yellow) and wild conditions during breeding (blue) and moulting (red).** Individual daily data points are represented in colour along with mean + SE in black.

**Table S1.** Parameter estimates for fixed effects in top ranked linear models for T_core_, T_eye_, T_bill_, T_flipper_, T_foot_ and T_trunk_ in relation to moult day. Time periods have been defined according to breakpoint analyses (see methods and results), except for non-moulting individual for which the entire study period has been considered. Significant p-values (<0.05) are denoted with (*).

| **Region** | **Time period (moult days)** | **Intercept (**°C) | **Moult day (**°C/day) | **p-value** | **T_w_ (**°C/°C_w_) or T_ground_ **(**°C/°C_ground_) | **p-value** | **R^2^_c_** |
| --- | --- | --- | --- | --- | --- | --- | --- |
| T_core_ | -28 to 3 | 38.48 | 0.064 | <0.0001* | NA | NA | 0.78 |
|  | 4 to 32 | 38.90 | -0.045 | <0.0001* | NA | NA | 0.70 |
|  | 34 to 62 | 37.35 | NA | NA | NA | NA | 0.26 |
|  | Non-moulting | 37.31 | NA | NA | NA | NA | 0.14 |
| T_eye_ | -37 to -10 | 26.36 | -0.08 | 0.0011* | NA | NA | 0.35 |
|  | -9 to 6 | 30.05 | 0.27 | <0.0001* | NA | NA | 0.63 |
|  | 7 to 27 | 32.92 | -0.17 | <0.0001* | NA | NA | 0.45 |
|  | 28 to 62 | 28.08 | NA | NA | NA | NA | 0.11 |
|  | Non-moulting | 25.05 | NA | NA | 0.34 | 0.0252* | 0.17 |
| T_bill_ | -37 to -12 | 15.34 | NA | NA | NA | NA | 0.24 |
|  | -11 to 6 | 17.85 | 0.68 | <0.0001* | 0.63 | 0.129 | 0.52 |
|  | 7 to 26 | 29.23 | -0.45 | <0.0001* | NA | NA | 0.33 |
|  | 27 to 62 | 16.08 | NA | NA | NA | NA | 0.30 |
|  | Non-moulting | 8.90 | NA | NA | 0.57 | 0.0058* | 0.12 |
| T_flipper_ | -37 to -11 | 9.43 | -0.08 | 0.0511 | NA | NA | 0.069 |
|  | -10 to -5 | 19.50 | 1.65 | 0.00775* | 1.36 | 0.05486 | 0.74 |
|  | -4 to 43 | 21.70 | -0.22 | <0.0001* | NA | NA | 0.54 |
|  | 45 to 62 | 4.09 | NA | NA | 0.87 | 0.00016* | 0.80 |
|  | Non-moulting | 7.21 | NA | NA | 0.54 | 0.0150* | 0.080 |
| T_foot_ | -37 to -10 | 11.31 | NA | NA | NA | NA | 0.47 |
|  | -9 to 6 | 9.37 | 0.21 | 0.12 | 0.25 | 0.014* | 0.65 |
|  | 7 to 13 | 23.35 | -0.77 | 0.04* | NA | NA | 0.44 |
|  | 15 to 62 | 13.59 | NA | NA | -0.087 | 0.0028* | 0.45 |
|  | Non-moulting | 7.89 | NA | NA | 0.069 | 0.0203* | 0.10 |
| T_trunk_ | -37 to -7 | 19.53 | NA | NA | -0.68 | 0.051 | 0.15 |
|  | -6 to 11 | 11.94 | -0.32 | <0.0001* | 0.29 | 0.054 | 0.55 |
|  | 12 to 25 | 5.68 | 0.42 | <0.0001* | NA | NA | 0.33 |
|  | 26 to 62 | 17.83 | -0.054 | 0.0153* | NA | NA | 0.55 |
|  | Non-moulting | 11.22 | NA | NA | 0.33 | 0.0097 | 0.093 |
